# Sex chromosome evolution, heterochiasmy and physiological QTL in the salmonid Brook Charr *Salvelinus fontinalis*

**DOI:** 10.1101/105411

**Authors:** Ben J. G. Sutherland, Ciro Rico, Céline Audet, Louis Bernatchez

**Affiliations:** Institut de Biologie Intégrative et des Systèmes (IBIS), Université Laval, Québec, QC, Canada G1V 0A6; School of Marine Studies, Molecular Diagnostics Laboratory, The University of the South Pacific, Laucala Campus, Suva, Fiji; Department of Wetland Ecology, Estación Biológica de Doñana (EBD-CSIC), c/Américo Vespucio s/n, 41092 Sevilla, Spain; Institut des Sciences de la Mer de Rimouski, Université du Québec à Rimouski, Rimouski, QC, Canada G5L 3A1

**Keywords:** heterochiasmy, salmon, sex chromosomes, QTL, whole genome duplication

## Abstract

Whole genome duplication can have large impacts on genome evolution, and much remains unknown about these impacts. This includes the mechanisms of coping with a duplicated sex determination system and whether this has an impact on increasing the diversity of sex determination mechanisms. Other impacts include sexual conflict, where alleles having different optimums in each sex can result in sequestration of genes into non-recombining sex chromosomes. Sex chromosome development itself may involve sex-specific recombination rate (i.e. heterochiasmy), which is also poorly understood. Family Salmonidae is a model system for these phenomena, having undergone autotetraploidization and subsequent rediploidization in most of the genome at the base of the lineage. The salmonid master sex determining gene is known, and many species have non-homologous sex chromosomes, putatively due to transposition of this gene. In this study, we identify the sex chromosome of Brook Charr *Salvelinus fontinalis* and compare sex chromosome identities across the lineage (eight species, four genera). Although non-homology is frequent, homologous sex chromosomes and other consistencies are present in distantly related species, indicating probable convergence on specific sex and neo-sex chromosomes. We also characterize strong heterochiasmy with 2.7-fold more crossovers in maternal than paternal haplotypes with paternal crossovers biased to chromosome ends. When considering only rediploidized chromosomes, the overall heterochiasmy trend remains, although with only 1.9-fold more recombination in the female than the male. Y chromosome crossovers are restricted to a single end of the chromosome, and this chromosome contains a large interspecific inversion, although its status between males and females remains unknown. Finally, we identify QTL for 21 unique growth, reproductive and stress-related phenotypes to improve knowledge of the genetic architecture of these traits important to aquaculture and evolution.

## INTRODUCTION

Characterizing the genetic architecture of ecologically-relevant phenotypes is essential for organism and genome evolution research (Rogers and Bernatchez 2007; Gagnaire *et al*. 2013) and selective breeding (Yáñez *et al*. 2014). Regions of the genome associated with specific traits can be identified by quantitative trait loci (QTL) analysis (Mackay 2001) or genome-wide association studies (GWAS; Bush and Moore 2012). Mapped traits can include morphological, behavioral, physiological or molecular phenotypes, but must show sufficient heritable variation. Power to detect QTL is determined by the number of individuals in the study (Slate 2013; Henning *et al*. 2014), the effect size of the QTL, allele frequencies (Mackay *et al*. 2009), the degree of polygenic control of the trait (Rockman 2012; Ashton *et al*. 2016), and the type of cross as well as the extent of the heritability of the trait (Mackay 2001). QTL mapping precision depends on recombination frequency (Mackay *et al*. 2009) as well as map density, although QTL are often only in linkage with causative mutations, which are rarely identified (Slate 2005). Trait genetic architecture can differ between families or populations (e.g. Santure *et al*. 2015) but parallelism and shared QTL can be identified (e.g. Laporte *et al*. 2015; Larson *et al*. 2015). It is therefore valuable to analyze multiple crosses to understand the broader implications of a QTL (e.g. Hecht *et al*. 2012; Palti *et al*. 2015; Lv *et al*. 2016). This can reduce false positives that may occur from small sample sizes (Slate 2013) by repeatedly identifying the same region associated to a trait, can more accurately determine the amount of variation explained by the QTL (Slate 2005), and can identify QTL that are not dependent on specific genetic backgrounds, which is particularly valuable for marker-assisted selection (Lv *et al*. 2016).

Advances in massively parallel sequencing (MPS) technology and comparative genomics have benefited QTL and association studies in several ways. MPS, or MPS-enabled technology such as high-density SNP chips greatly increases marker density (Catchen *et al*. 2011; Ashton *et al*. 2016), provides flanking sequence of markers that can be indexed in reference to or aligned against reference genomes for integrating across species (Sutherland *et al*. 2016) or for identifying genes near QTL to inform on potential drivers underlying a trait (e.g. McKinney *et al*. 2016; Johnston *et al*. 2016). When causative mutations are not known, detecting orthologous QTL in other species can provide further evidence for a region or gene being related to a trait (Mackay 2001; Larson *et al*. 2015). As an example, QTL for recombination rate in mammalian model and non-model systems occur near the same genes (Johnston *et al*. 2016). Comparative genomics clearly has an important role in identifying drivers of trait variation.

Genetic architecture can be strongly affected by sexually antagonistic selection and sex determination. Sexually antagonistic alleles (i.e. alleles that benefit sexes differently) produce genetic conflict (Mackay 2001; Charlesworth *et al*. 2005), which can be resolved by the sequestration of alleles in non-recombining sex chromosomes. As an example of the effect this can have on genome architecture, the *Drosophila* Y chromosome is made almost exclusively of genes that have migrated from other chromosomes, presumably due to their specific benefit to males (Carvalho 2002). An additional benefit occurs by constant sex-specific selection occurring for alleles on the Y (or W) chromosome, as these are always only in the heterogametic sex (Lahn *et al*. 2001). However, the lack of recombination between the sex chromosomes can also result in Y degeneration due to accumulation of mutations that are not able to be purged through recombination with X (Charlesworth 1991). A different resolution to genetic conflict involves sex-dependent dominance, whereby allelic dominance depends on the sex of the individual, which need not be on the sex chromosome (Barson *et al*. 2015). Much remains to be understood about resolving these conflicts.

Genome evolution can also be affected by large mutational forces, such as polyploidization events including whole genome duplication (WGD) (Ohno 1970), which may disrupt sex determination systems (Davidson *et al*. 2009). Although details of this disruption remain generally unknown, some hypotheses have been proposed involving the independent segregation of duplicated sex determining chromosomes or imbalances in gene dosages when X inactivation occurs (Muller 1925; Orr 1990; Davidson *et al*. 2009). Highly diverse sex determination systems are observed in teleosts (Marshall Graves and Peichel 2010). This diversity may have been influenced by the teleost-specific WGD due to a post-WGD adoption of numerous different sex determination mechanisms (Mank and Avise 2009). Evolution of sex determination post-WGD may occur through the mutational disruption of one duplicated portion of the existing system or by the development of a new system (Davidson *et al*. 2009), which can thus result in the evolution of new sex chromosomes.

Sex chromosome evolution may be facilitated by differences in recombination rates between the sexes (i.e. heterochiasmy) (Charlesworth *et al*. 2005), where tight linkage forms along the sex chromosome in the heterogametic sex (Haldane 1922). The evolution of heterochiasmy remains under investigation, although several explanations have been proposed (Lenormand and Dutheil 2005; Brandvain and Coop 2012; Lenormand *et al*. 2016). First, sexes can experience different extents of selection at haploid stages (Lenormand 2003), and heterochiasmy permits retention of epistatically-interacting alleles within a haplotype specifically within the sex experiencing more haploid selection (Lenormand and Dutheil 2005). Second, physical meiotic differences may play a role; female meiosis occurs with a long delay, and chiasma (i.e. locations where crossovers occur) stabilize chromatids during this process (Lenormand 2003; Lenormand *et al*. 2016). Third, recombination protects from meiotic drive, to which the sexes have different susceptibilities (Brandvain and Coop 2012; Johnston *et al*. 2017). Other hypotheses have also been proposed (Trivers 1998; Lenormand 2003). However, in general it is unclear which of the above explanations have the largest influence, and thus the relationships between WGD, heterochiasmy and sex chromosome evolution require further study.

Salmonids (Family Salmonidae) are an ideal system to study genetic architecture and sex determination post-WGD (Davidson *et al*. 2010). The salmonid genome remains in a residually tetraploid state, where some chromosomal telomeric regions continue recombining between homeologous chromosomes and others have rediploidized (Allendorf and Thorgaard 1984; Allendorf *et al*. 2015; May and Delany 2015; Lien *et al*. 2016). Salmonid sex determination is genetically controlled (Davidson *et al*. 2009) by a truncated gene from the *interferon-response factor* transcription factor family, *sdY* (sexually dimorphic on the Y-chromosome; Yano *et al*. 2012a). *sdY* may be a salmonid innovation as it has not yet been identified in the non-duplicated sister group for the salmonid WGD, Northern Pike *Esox lucius* (Yano *et al*. 2012b). Male genome-specific conservation of *sdY* occurs in more than ten salmonid species, but some exceptions exist, including the Lake Whitefish *Coregonus clupeaformis* and European Whitefish *C. lavaretus* (Yano *et al*. 2012b), and some Atlantic Salmon *Salmo salar* and Sockeye Salmon *Oncorhynchus nerka* individuals (Eisbrenner *et al*. 2013; Larson *et al*. 2016). Sex chromosomes are not homologous among many salmonid species, potentially due to transposition of *sdY* between chromosomes (Woram *et al*. 2003). Additional evidence for transposition includes repetitive flanking regions with putative transposable elements (Brunelli *et al*. 2008) (Lubieniecki *et al*. 2015) and sequence conservation that abruptly stops outside of the sex determination cassette (Faber-Hammond *et al*. 2015). This transposition to different chromosomes may be delaying Y degeneration (Yano *et al*. 2012b; Lubieniecki *et al*. 2015). In general, the salmonids are at an early stage of sex chromosome evolution (Phillips and Ihssen 1985; Yano *et al*. 2012b) where sex chromosomes are homomorphic (Devlin *et al*. 1998; Phillips and Ráb 2001; Davidson *et al*. 2009). Male salmonids have low recombination rates relative to females with crossover events primarily occurring at telomeric regions, as observed in Rainbow Trout *O. mykiss* (Sakamoto *et al*. 2000) and Atlantic Salmon (Moen *et al*. 2004). Heterochiasmy was principally observed in one linkage group in Northern Pike (i.e. the sister species of the salmonid WGD), but in general was much more equal between the sexes than previously observed in salmonids (Rondeau *et al*. 2014). Salmonids are therefore a valuable model to study the evolution and effects of heterochiasmy in relation to sex determination post-WGD.

The combination of characterizing heterochiasmy, sex chromosome identity and the genetic architecture for reproductive, growth and stress response traits provides much-needed information regarding the function of the Brook Charr *Salvelinus fontinalis* genome post-duplication. The goals of this study are to use a high-density genetic map for Brook Charr (Sutherland *et al*. 2016) to (*a*) identify the sex-linked chromosome; (*b*) quantify heterochiasmy in this mapping family while correcting for probable genotyping errors; and (*c*) search for growth, stress resistance and reproduction-related QTL. Furthermore, using the recent characterization of homology to ancestral chromosomes and homeolog identification among the salmonids (Sutherland *et al*. 2016), we subsequently compare identities of sex chromosomes and identified QTL across the salmonids to identify consistencies. We then discuss the implications of sex chromosome consistencies and heterochiasmy in relation to sex chromosome evolution in salmonids.

## METHODS

### Fish and phenotyping

Juvenile Brook Charr used in this study were the same individuals used to construct a low-density genetic map and perform QTL analysis for 21 phenotypes (29 including repeated measurements occurring at three time points; Table S1) by Sauvage *et al*. for growth (2012a) and reproductive QTL (2012b), as well as to produce a high-density genetic map (Sutherland *et al*. 2016). In brief, F_0_ individuals were from a wild anadromous population and a domestic population, and two of the F_1_ individuals were crossed to produce 192 F_2_ offspring. After filters for missing data per individual, 22 offspring were excluded, and one was excluded due to abnormally high numbers of crossovers, leaving 169 individuals (Sutherland *et al*. 2016). Fish were raised in tanks as previously described until 65-80 g, at which point weight, length and condition factor were measured. These phenotypes were measured on the same fish two and six months after the initial measurements. Growth rate was calculated between the multiple sampling times. At the final sampling, all phenotypes were collected. Stress response was also evaluated at this final sampling through an acute handling stress by reducing water levels, capturing fish without chasing and holding out of water for one minute in order to phenotype the stress response using blood parameters chloride, osmolality and cortisol before and after the stress. After fish had re-acclimatized, they were anaesthetized and killed by decapitation as per regulations of Canadian Council of Animal Protection recommendations and protocols approved by the University Animal Care Committee, as previously reported (Sauvage *et al*. 2012a). The sex of each individual was determined by visual inspection of the gonads as reported by Sauvage *et al*. (2012b).

### Genetic map and quality control of markers and phenotypes

A recently developed high-density genetic map with 3826 markers was used with genotypes for 192 offspring (Sutherland *et al*. 2016). Parents were diploid and therefore the map is probably missing residually tetraploid regions because these would be removed due to too many alleles during genotyping (see Sutherland *et al*. 2016 for more information). In brief, genotype data was obtained using the population module of STACKS v.1.32 (Catchen *et al*. 2011), phased in JoinMap v.4.1 (van Ooijen 2006), and imported into R/qtl (Broman *et al*. 2003) using the *read.cross* function with data interpreted as a four-way cross type in the *mapqtl* format (see File S1 for map, genotype and phenotype input files).

All 29 phenotypes (including eight measures at multiple time points) related to blood parameters, growth, growth-related gene expression, reproduction and stress response were used to search for QTL (Table S1). Correlation between phenotypes was evaluated using Pearson correlation in R (R Development Core Team 2017) and a correlation plot was generated using the R package *corrplot* (v.0.77; Wei and Simko 2017). Phenotypes were inspected for normal distribution, and when required, log transformed (Broman and Sen 2009). Outlier phenotype values (>3 SD from the mean) were removed to prevent spurious associations (Broman and Sen 2009), including two individuals each for T1-T2 and T2-T3 growth rates, two individuals for length at T2, four individuals for condition factor at T2, one individual for change in osmolality and one individual for sperm diameter.

Markers present in the map were tested for segregation distortion by chi-square tests for Mendelian segregation in R/qtl and removed when p ≤ 0.01 (Broman and Sen 2009). A total of 157 markers with significant segregation distortion were removed, leaving a remainder of 3669 markers. Proportions of identical genotypes were tested in R/qtl to ensure that there were no mis-labeled samples. Recombination fraction between marker pairs was estimated using Expectation Maximization algorithm within *est.rf* in R/qtl. The minimum number of obligate crossover events was calculated per individual using *count.XO* in R/qtl, and an outlier sample with 1093 crossovers was removed (other samples had mean and median crossovers of 101 and 83, respectively, before correcting for unlikely double crossovers).

### Recombination rate

To characterize heterochiasmy in the mapping family parents, the *plotGenotypes* function of R/qtl was used to identify positions of crossovers per parental chromosome (total = 84 chromosomes per individual offspring) and modified to export these positions (see Data Availability section for all code used in the analysis). Male-specific markers were not included in the original map due to low recombination rate and poor positioning (Sutherland *et al*. 2016), and therefore to avoid bias of including female-specific but not male-specific markers, crossovers were evaluated in a map with only markers informative in both sexes (i.e. *ef* x *eg* and *hk* x *hk*; see van Ooijen 2006 and Wu *et al*. 2002 for full explanation of marker types for a cp cross type). Furthermore, as recombination rates can be inflated by a genotyping error appearing to be flanked by two false recombination events (Hackett and Broadfoot 2003; Slate 2008), which can also occur in RAD-seq data (Andrews *et al*. 2016), an additional correction was made to more accurately quantify heterochiasmy. Specifically, per individual and per phased haplotype within individual, whenever a crossover is identified, a search within 50 cM of the crossover is conducted to identify if a second crossover is near (or an even total of crossovers, resulting in no true phase change). If this number is even, it suggests that the putative crossovers may have been due to a genotyping error, as double crossovers are not expected due to crossover interference. As such, these crossovers would not be counted in the total sum. This is similar to the approach used by Johnston *et al*. (2016) to avoid false double crossovers by only including crossovers that flank more than a single marker. Subsequently, the cumulative number of crossovers for fused metacentric and acrocentric chromosome were calculated and cumulatively displayed in positions as a percentage of the total chromosome length. The corrected crossover counts were used to calculate the female:male recombination rates of the parents. This was also conducted without cumulating and displayed on a per chromosome per haplotype basis.

### QTL analysis

The effect of sex on each phenotype was tested using linear models in R (R Development Core Team 2017). If a marginal effect of sex was found (p ≤ 0.20), sex was included in the model as a covariate for the phenotype to reduce residual variation and improve power to identify the QTL (Broman and Sen 2009). The R/qtl function *scanone* with permutation testing (10,000 permutations; p ≤ 0.05) was used to identify the presence of a single QTL within each linkage group (Broman *et al*. 2003). Chromosome-wide significance was tested in the same way but per chromosome (10,000 permutations; p ≤ 0.01). Confidence interval estimates (95%) for QTL positions were identified using *summary.scanone* calculating LOD support intervals with a 1.5 LOD drop. Sex-specific phenotypes (i.e. sperm diameter and concentration, egg diameter) were tested in only one sex, and therefore had smaller sample sizes. Percent variance explained by the identified QTL was performed using *makeqtl* and *fitqtl* within R/qtl, including all genome and chromosome-wide QTL per trait in the formula (trait ∼ QTL_1_ + QTL_2_ + QTL_n_), as well as sex as a covariate when required. Phenotypic effects were estimated by calculating the differences between the mean phenotype values among the genotype groups for the marker closest to the identified QTL, including only individuals that were successfully genotyped. For markers that only segregate in one parent (i.e. *nn* x *np*) only two phenotype by genotype averages are given, one for the homozygote and one for the heterozygote offspring. Alternatively, for markers segregating in both parents (i.e. *hk* x *hk* or *ef* x *eg*), three phenotype averages are given, two for the alternate homozygotes and one for the heterozygote in *hk* x *hk* marker types and two for the alternate heterozygotes and one for the homozygote in *ef* x *eg* marker types. Sex-specific averages were calculated when the QTL required sex as a covariate in the model. RAD tags for all alleles and associated QTL results are available in File S2.

To identify the sex chromosome, offspring sex was coded as a binary trait to identify linkage to any of the LG by QTL mapping as described above (Broman and Sen 2009). Furthermore, the effect of a QTL may vary depending on the sex of an individual in a non-additive manner (Broman and Sen 2009) due to genetic variation in sexual dimorphism for the trait (e.g. loci that have a different effect in males and females Mackay 2001). Therefore, QTL by sex interaction effects were inspected per trait by subtracting an additive model (*genotype* and *sex*) from a full model (*genotype*, *sex* and a *sex-by-genotype* interaction term) as described by Broman and Sen (2009). If the additive model is largely driving the effect, the model with only the interaction effect will not be significant, and in this case the interaction effect would not be included in the model. As suggested by Broman and Sen (2009), interaction effects were only tested in this way when a full model for a locus was significant.

Identities of sex chromosomes of other species were obtained from references listed in Table 1. For Atlantic Salmon, Artieri *et al*. (2006) identify that the sex determining region is on the long (q) arm of chromosome Ssa02, and Lien *et al*. identify that Ssa02q is homeologous to Ssa12q, indicating that the chromosome arm holding the sex determining region corresponds to the ancestral chromosome 9.1 (Sutherland *et al*. 2016). Other species were directly obtained from references in Table 1. Correspondence between Arctic Charr *S. alpinus* and Brook Charr were identified indirectly through other species shared between Nugent *et al*. (2016) and Sutherland *et al*. (2016).

**Table 1.**
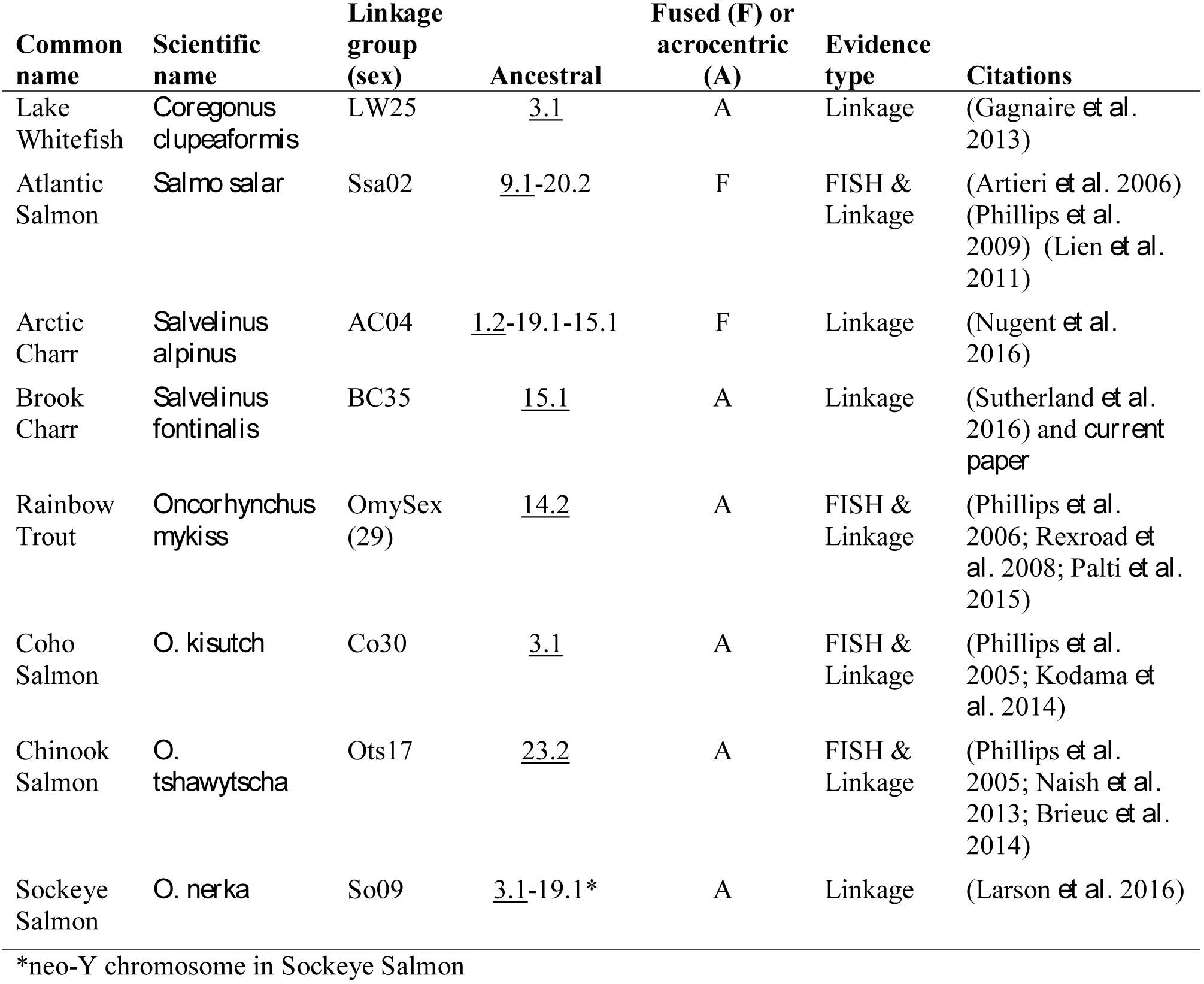
Salmonid sex chromosomes from high-density genetic maps named with Northern Pike designations (ancestral). The chromosome arm that contains the sex determining region is underlined, and the fusion status of the chromosome and original reference are provided. Ancestral chromosomes are defined by Sutherland *et al*. (2016) and are based on Northern Pike chromosomes from Rondeau *et al*. (2014).

### Data availability

The raw data for this study is available in the NCBI SRA in BioProject PRJNA308100 and accession SRP068206. All input files used for the analysis are in the supplementary files (File S1) and all code used to perform analyses is available on Github at the following link: https://github.com/bensutherland/sfon_pqtl/

## RESULTS

### Sex-linked chromosome in Brook Charr

Sex was highly associated with the majority of Brook Charr (BC) linkage group (LG) BC35, indicating that this is the sex-linked chromosome in Brook Charr (Figure 1). Linkage across the entire LG until the LOD score decreases at the distal end can be explained by male salmonid-specific low recombination rate and male bias of crossovers towards telomeric regions (Sakamoto *et al*. 2000). The drop in LOD suggests the far end of the chromosome is pseudoautosomal, which even occurs in the highly differentiated mammalian X/Y chromosomes in a recombinogenic distal region of the Y chromosome (Lahn *et al*. 2001). Many of the recombination events in BC35 were at a similar section of the LG (∼90-110 cM; Figure S1B).

**Figure 1.**
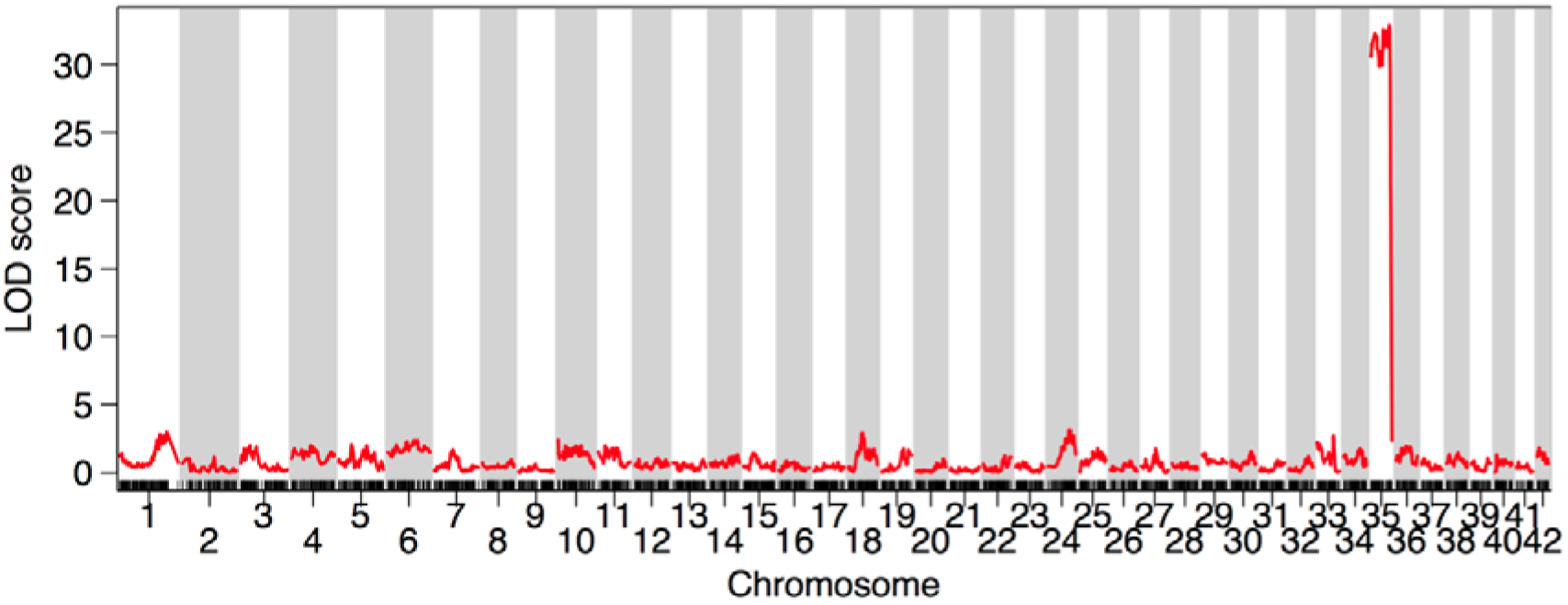
The acrocentric linkage group BC35 is highly associated with sex in Brook Charr. Due to low recombination in males, high linkage is viewed across the majority of the linkage group.

BC35 is an acrocentric chromosome homologous to the Northern Pike chromosome 15.1 (homeolog naming from Sutherland *et al*. 2016). There are no other sex chromosomes in the other salmonid species with high-density genetic maps available with the chromosome arm containing the sex determining region homologous to BC35 (Table 1). Arctic Charr has a sex chromosome that is comprised of a triple fused chromosome (although this may vary across populations) that contains 15.1 in the fusion, but in Arctic Charr this is not the chromosome arm that holds the sex determining region, which is held within the arm on the other side of the chromosome AC04p (Nugent *et al*. 2016). Other salmonids have different sex chromosomes, as shown in Table 1, including Lake Whitefish (3.1; Gagnaire *et al*. 2013), Atlantic Salmon (9.1-20.2; Artieri *et al*. 2006; Lien *et al*. 2011), Rainbow Trout (14.2; Palti *et al*. 2015), Coho Salmon *O. kisutch* (3.1; Phillips *et al*. 2005; Kodama *et al*. 2014), Chinook Salmon *O. tshawytscha* (23.2; Phillips *et al*. 2005; Naish *et al*. 2013; Brieuc *et al*. 2014) or Sockeye Salmon (3.1-19.1; Larson *et al*. 2016). This further refines previous observations of the general lack of homology in the sex chromosomes of the salmonids (Woram *et al*. 2003). Some information on sex chromosomes identities across *Salmo, Salvelinus* and *Oncorhynchus* have been previously reported (Phillips 2013) and most of the results correspond with those here, with the exception of the Brook Charr sex chromosome, which the two studies identify as corresponding to opposite arms of the Arctic charr sex chromosome. This i possibly due to a population polymorphism, but more work would be needed to confirm this.

Considering the importance of inversions to sex chromosome formation through the reduction of recombination between X and Y (Lahn *et al*. 2001; van Doorn and Kirkpatrick 2007; Berset-Brandli *et al*. 2008), it is interesting to note that Brook Charr has a species-specific inversion in BC35 in the female map (15.1; see Figure 5 in Sutherland *et al*. 2016). As is usual for salmonid linkage maps, the male-specific map was not produced as the low recombination frequency resulted in poorly placed male-specific markers (Sutherland *et al*. 2016), and so it is not possible to check whether this inversion is heterozygous within the species, but this will be valuable to investigate in future studies.

### Sex-specific recombination rate and positions of crossovers

Crossovers occurred 2.7-fold more often in the maternal haplotypes (total = 3679) than in the paternal (total = 1368; Figure 2) based on the phased haplotypes of 169 individual offspring (Wu *et al*. 2002; Sutherland *et al*. 2016). The double recombinant correction (see Methods) in the autosomes removed 606 and 682 crossover events due to probable genotyping errors from the dam and sire, respectively, providing a more accurate estimation of the heterochiasmy ratio, although the trends regarding the crossover positions remained similar. Crossovers were biased towards the center of the linkage groups in the dam and towards the external 20% of the linkage groups in the sire (Figure 2). This bias is similar to that observed in Rainbow Trout (Sakamoto *et al*. 2000), and although reasons for it remain under investigation, there are currently several hypotheses that may explain it including protection against selfish genetic elements and meiotic drive in female meiosis (Brandvain and Coop 2012; Johnston *et al*. 2017), among others (see Introduction). When the distance between putative recombination events was reduced to 25 cM or 10 cM for the double-recombinant correction method, the overall trend of more recombination in the female remained (2.5 and 2.2-fold more recombination in the female, respectively), and the spatial bias with male recombination near telomeres and female throughout the chromosome remained.

**Figure 2.**
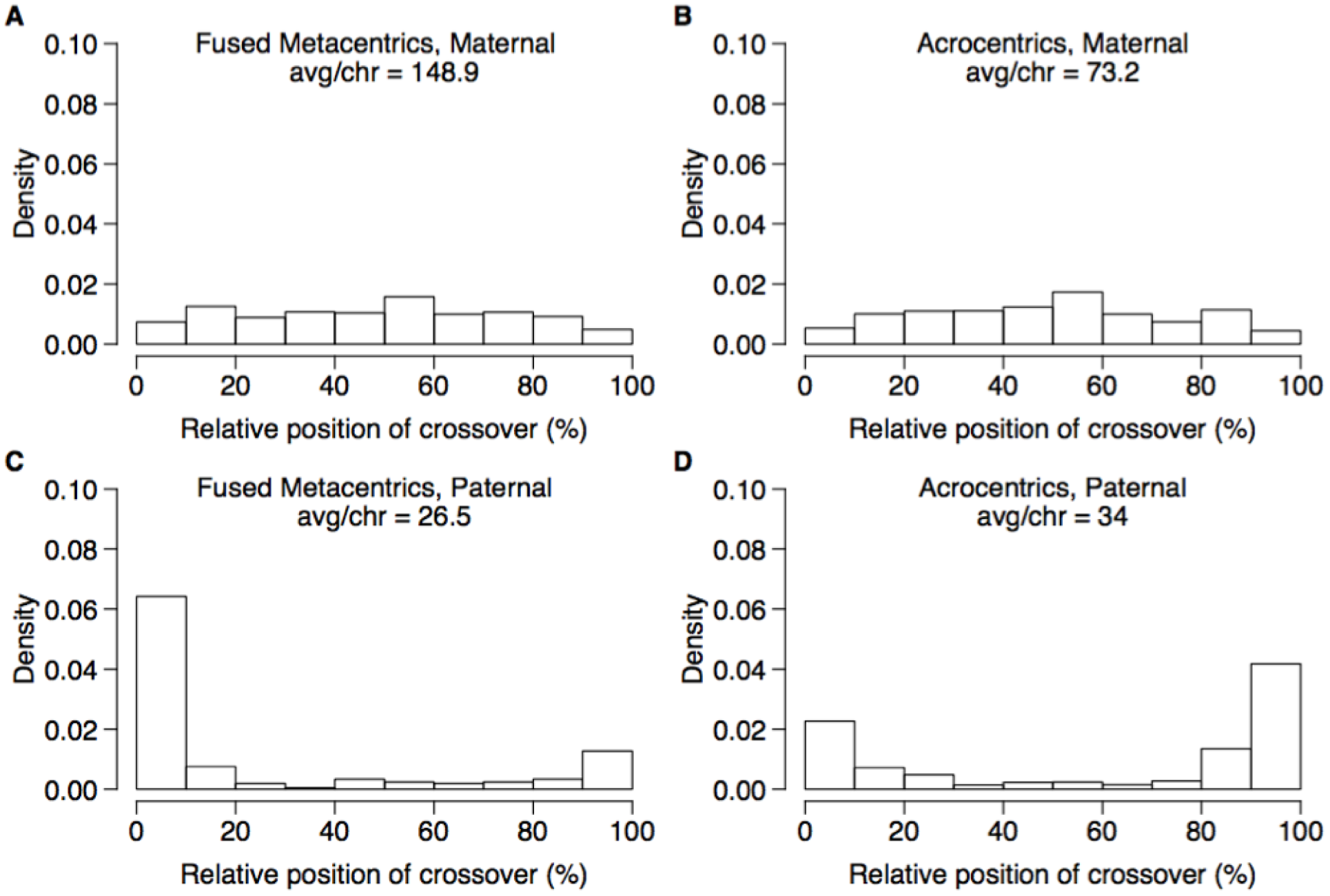
Maternal and paternal cumulative crossover positions across the chromosomes. The position of each crossover is expressed as a percent of the total crossover length and cumulated for all crossovers within each chromosome type, specifically fused metacentric chromosomes (A,C) and acrocentric chromosomes (B,D) in the maternal and paternal haplotypes, respectively. Maternal haplotypes had 2.7-fold more crossovers than paternal haplotypes, with the maternal crossovers occurring throughout the chromosome and the paternal crossovers restricted mainly to the first and/or last 20% of the linkag groups.

Separating chromosomes into fused metacentric (n = 8) and acrocentric chromosomes (n = 34) indicated a higher heterochiasmy ratio in fused metacentric than acrocentric chromosomes (5.6-fold and 2.2-fold, respectively). The male had fewer crossovers per chromosome in the fused metacentrics (mean = 26.5) than acrocentrics (mean = 32.9), even though fused metacentrics are comprised of two acrocentric chromosomes combined and thus are longer. In contrast, the female had approximately twice as many crossovers per fused metacentric chromosome (mean = 148.9) than acrocentric (mean = 70.5). The lower recombination in the paternal fused metacentrics than the paternal acrocentrics is probably due to missing regions of the genetic map that are residually tetraploid that were removed during marker filtering due to quality filtering, as the map was produced using a diploid cross (Limborg *et al*. 2016). Inspection of individual chromosomes indicates that the chromosomes expected to still exhibit residual tetraploidy (Sutherland *et al*. 2016) all show a lack of crossovers in the male relative to the chromosomes expected to have returned to a diploid state (Figure S1). The missing regions in the 16 (of 50) chromosome arms expected to be residually tetraploid will result in an inflation of the heterochiasmy ratio, as male crossovers will be specifically underestimated for these arms. This was confirmed by separating chromosomes into those expected to exhibit residual tetraploidy or to be rediploidized (Sutherland *et al*. 2016) and recalculating the heterochiasmy ratio. For residually tetraploid chromosomes, including both arms when present in a fused chromosome (total = 16 chromosomes), the female:male heterochiasmy ratio is 6.4 (female = 1608; male = 251) and for expected rediploidized chromosomes (total = 26) the ratio is 1.85. The overall ratio is therefore probably inflated due to missing regions of the male map, and when the residually tetraploid chromosomes are removed, the ratio remains female-biased at 1.85-fold more than male. Furthermore, biased positions of crossovers remain regardless of residual tetraploid status where the female is towards the center and the male towards the external sides of the chromosome (Figure S1). In summary, the male has fewer recombination events than the female and the crossovers are biased to the distal portions of the chromosomes.

The identified sex chromosome had more crossovers than the average male acrocentric chromosomes (sex = 71, other acrocentrics average = 32.9), and all of the crossover events in the sex chromosome occurred at one end of the chromosome and almost no crossovers occurred in the rest of the chromosome (Figure S1). This bias to only a single end of each chromosome in the male map was consistent throughout all of the chromosomes with crossovers present. Using the positions of centromeres determined for Chinook Salmon (Brieuc *et al*. 2014), and placing them in the corresponding position on the Brook Charr map using map correspondence (Sutherland *et al*. 2016), indicates that the end of the acrocentric chromosomes where the crossovers occur is the opposite end to that containing the probable centromere (see Figure S1). Using this information, the few occurrences of unknown centromere positions can be easily determined as they are probably at the opposite side to where the crossovers occur.

### QTL identification: growth, reproduction and stress response

Genome-wide significant QTL were identified for weight, length, condition factor, specific growth rate, and liver weight (Table 2; Table S2). A total of 29 QTL were found to be significant at the chromosome-wide level (p ≤ 0.01), and these included QTL for phenotypes egg and sperm diameter, change in cortisol, chloride and osmolality after an acute handling stress, *growth hormone receptor* gene expression and hematocrit (Table 2). In total, QTL were identified on 14 of the 42 Brook Charr linkage groups (Figure 3).

**Figure 3.**
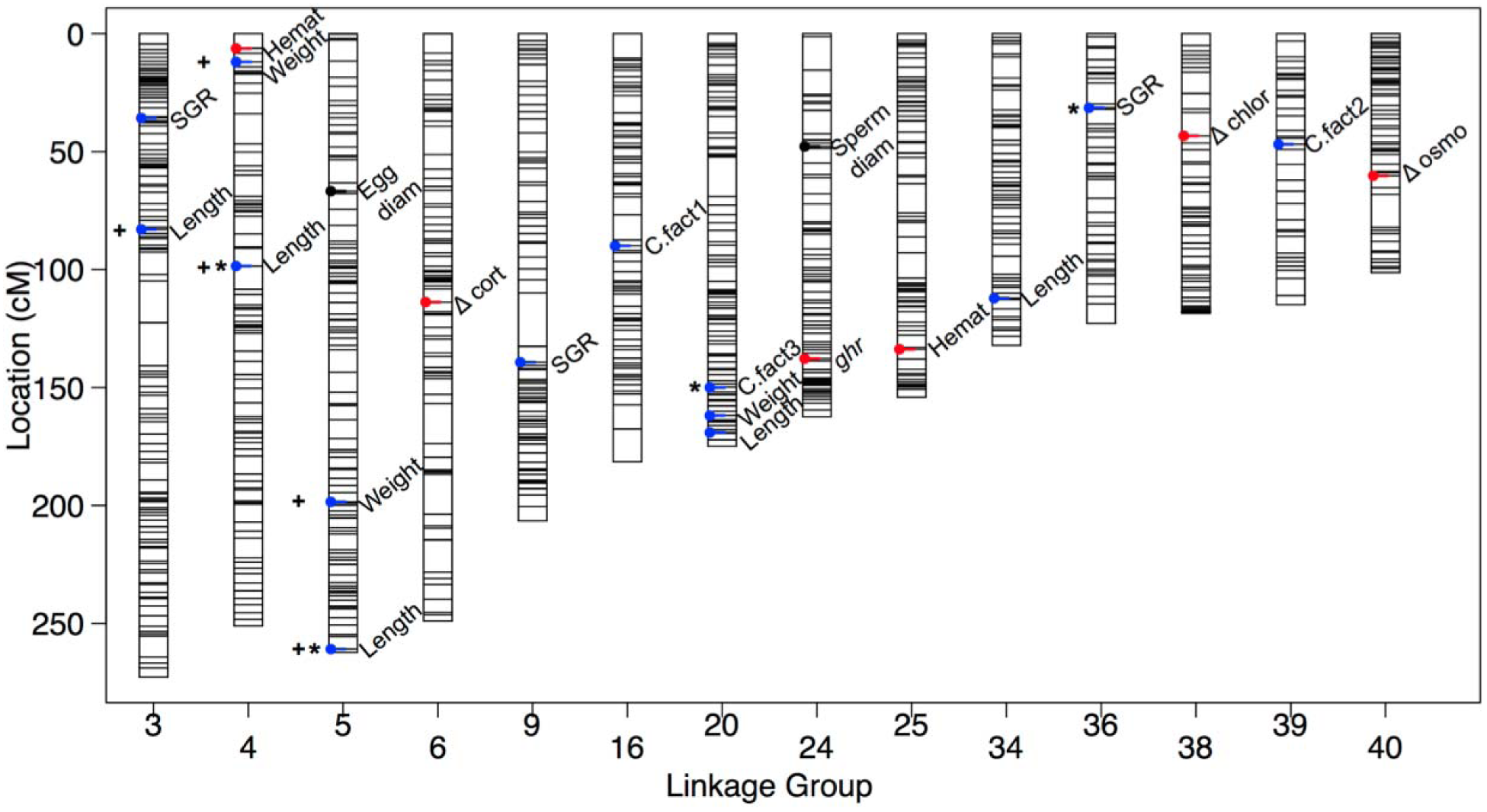
All identified QTL plotted on the Brook Charr genetic map. QTL for growth related traits are shown in blue, reproductive in black, and blood or stress-related in red. QTL with asterisks are at the genome-wide significance level, and the rest are chromosome-wide. QTL with broad confidence intervals discussed in the Results are denoted with a positive symbol (+). More details on phenotypes can be found in Table S1 and on QTL can be found in Table 2 and Table S2.

**Table 2.** Identified QTL in Brook Charr with positions, percent variance explained (PVE) and the effect of the allelic state on the trait. Sex was included as a covariate when required, and in these cases the allelic effect is given for both males and females, and the PVE from sex is also given. When sex was not required as a covariate, the second averages are displayed as NA and the first averages represent both sexes. The phenotype average for the homozygote common allele (aa avg) is shown for comparison to the largest effect size (effect ♀ or ♂). LG = linkage group; Pos = cM position; CI = confidence interval; PVE = percent variance explained; sex.cov = sex included as a covariate. QTL significance is displayed as genome wide p ≤ 0.01 ***; p ≤ 0.05 **; or chromosome-wide p ≤ 0.01 *.

| Phenotype | LG | Pos | 95% CI | Marker | QTL pval | Tot. PVE | Ind. PVE | aa avg ♀ | effect ♀ | aa avg ♂ | effect ♂ |
| --- | --- | --- | --- | --- | --- | --- | --- | --- | --- | --- | --- |
| Weight (g) T2 | 4 | 28.3 | 16-214 | 7187 | * | 42.8 | 9.5 | 126 | +22.1 | 129.5 | +52.1 |
|  | 5 | 198 | 39-262 | 125487 | * |  | 8.1 | 121.4 | +27.9 | 145.3 | +30.6 |
|  | 20 | 162 | 105-175 | 6352 | * |  | 6.2 | 133.9 | -2.2 | 171.2 | -22.8 |
|  |  |  |  | sex.cov |  |  | 9.5 |  |  |  |  |
| Length (cm) T2 | 3 | 83 | 19-267 | 90770 | * | 54.5 | 2.0 | 22.4 | -0.3 | 23.6 | +0.4 |
|  | 4 | 115 | 16-215 | 66075 | ** |  | 5.5 | 21.8 | +0.9 | 23.6 | +0.2 |
|  | 5 | 261 | 185-262 | 85980 | *** |  | 8.2 | 21.7 | +1.1 | 23.6 | +0.3 |
|  | 20 | 169 | 99-175 | 60142 | * |  | 4.1 | 21.7 | +0.8 | 22.9 | +1.3 |
|  | 34 | 112 | 85-132 | 120757 | * |  | 3.1 | 22.5 | -1.1 | 24.0 | +0.7 |
|  |  |  |  | sex.cov |  |  | 13.5 |  |  |  |  |
| Cond. Fact. T1 | 16 | 89.9 | 48-105 | 118085 | * | 10.0 | 10.0 | 1.0 | -0.02 | NA | NA |
| Cond. Fact. T2 | 39 | 46.9 | 35-83 | 39977 | * | 10.3 | 10.3 | 1.2 | -0.03 | NA | NA |
| Cond. Fact. T3 | 20 | 150 | 116-175 | 55565 | ** | 12.2 | 12.2 | 1.1 | +0.04 | NA | NA |
| SGR T2-T3 | 3 | 35.7 | 18-85 | 115199 | * | 26.0 | 6.8 | 0.6 | +0.10 | NA | NA |
|  | 9 | 139 | 89-189 | 128240 | * |  | 5.3 | 0.6 | +0.04 | NA | NA |
|  | 36 | 31.4 | 1-80 | 30493 | *** |  | 5.3 | 0.6 | -0.09 | NA | NA |
| Egg diameter | 5 | 66.8 | 41.7-185 | 37572 | * | 39.0 | 38.95 | 4.0 | -0.046 | NA | NA |
| Sperm diameter | 24 | 47.8 | 0-59.8 | 202134 | * | 32.6 | 32.59 | NA | NA | 2.9 | -0.001 |
| Δ cortisol | 6 | 114 | 108-135 | 113752 | * | 57.6 | 9.0 | 7.4 | +2.3 | 0.8 | -0.6 |
|  |  |  |  | sex.cov |  |  | 43.0 |  |  |  |  |
| Δ chloride | 38 | 43.3 | 25.3-60.9 | 116693 | * | 18.5 | 18.5 | 0.3 | -2.9 | NA | NA |
| Δ osmolality | 40 | 60.3 | 26.3-82.1 | 52306 | * | 31.1 | 14.0 | 15.2 | -3.4 | -5.4 | +4.6 |
|  |  |  |  | sex.cov |  |  | 14.6 |  |  |  |  |
| *ghr* | 24 | 138 | 47.8-153 | 141355 | * | 23.8 | 23.8 | -3.9 | +0.2 | NA | NA |
| Hematocrit | 4 | 22.6 | 16.3-161 | 105237 | * | 42.8 | 12.0 | 35.3 | +0.9 | 39.6 | -1.7 |
|  | 25 | 139 | 113-159 | 1153 | * |  | 12.4 | 35.1 | +0.5 | 38.3 | -0.2 |
|  |  |  |  | sex.cov |  |  | 13.9 |  |  |  |  |

Several traits showed sexual dimorphism and therefore required sex as a model covariate. These included weight, length, liver weight, hepatosomatic index, hematocrit, change in osmolality and cortisol from stressor, resting plasma chloride, hepatic glycogen, *insulin-like growth factor 1* and *igf receptor 1* (Table S1). Specific growth rate, condition factor, change in chloride, resting plasma osmolality and glucose, and *growth hormone receptor* gene expression did not show sexual dimorphism. Traits with high phenotypic correlation included length and weight (r = 0.90 at T1), and liver weight and hepatosomatic index (r = 0.85; Figure S2). Specific growth rate T1-T2 was negatively correlated with weight at T1 (r = - 0.64), suggesting that larger individuals measured at T1 subsequently grew slower than smaller individuals. Other traits generally were not as highly correlated (r < 0.35). Even though the phenotypes were not highly correlated, QTL were identified for condition factor and weight in the same region of BC20, and QTL affecting hematocrit and weight (r = 0.24) were found in the same region on BC04 (Figure 3).

A few specific trait-linkage group combinations had elevated LOD across a large portion of the LG. This was observed for length and weight on BC03, BC04 and BC05. To determine if this was due to one specific marker type, the three marker types were each tested independently for QTL (female-specific *nn* x *np*; informative in both parents *ef* x *eg*; and semi-informative *hk* x *hk*). Interestingly, when including markers only polymorphic in the female (i.e. *nn* x *np*) these elevated LOD baselines were not observed (*data not shown*). It is possible that this may be therefore due to an effect originating from paternal alleles, which have strong linkage across the entire LG. As these QTL explained a substantial amount of variance for these three traits, the full analysis includes these markers, but the exact locations within the LG of these QTL cannot be determined without additional families or crosses (markers noted in Figure 3).

A substantial amount of trait variance for length at T2 within this mapping family was explained by five QTL. Together with the additive sex covariate, this collectively explained 54.5% of the trait variation. The QTL with broad elevated LOD on BC04 and BC05 (see above) individually explained 5.5 and 8.2% of the variation, respectively. QTL for condition factor varied depending on the sampling time (T1-T3), where each time point had a QTL at a different LG that explained over 10% of the trait variation within the time point. However, trait variation was small for this trait and therefore effect sizes of the QTL were also small (Table 2). QTL for specific growth rate (SGR) were identified on BC03, BC09 and BC36, with three QTL explaining 26% of the SGR (T2-T3) trait variation (Table 2).

Reproductive traits were sex-specific and therefore had approximately half of the individuals as the traits with values in both sexes, and thus had less statistical power, reducing the ability to detect QTL. Nonetheless, enough power was present in the data to detect chromosome-wide significant QTL for egg and sperm diameter (Table 2; Figure 3). A QTL for egg diameter was identified at 67 cM of BC05, explaining 39% of the trait variation and a QTL for sperm diameter was identified at 48 cM of BC24, explaining 33% of the trait variation. Neither of these reproductive-related traits mapped to the sex-linked chromosome (BC35). A QTL for *growth hormone receptor* (*ghr*) gene expression was identified at 138 cM of BC24 explaining 24% of the trait variation.

Stress response QTL were identified only at the chromosome-wide significance level. This included responses of cortisol (114 cM of BC06), chloride (43 cM of BC38), and osmolality (60 cM on BC40; Table 2; Figure 3). Change in cortisol from acute handling stress was highly dependent on sex; the identified QTL explained 9% of the trait variation whereas sex explained 43% (total PVE = 58%; Table 2). Females heterozygous at the marker closest to the QTL increased cortisol by 2.3 μg/dL plasma more than the homozygote, and heterozygous males had 0.65 μg/dL lower than the homozygote. To further demonstrate the large sex effect, averaging the two genotypes shows that females increased blood cortisol by 8.6 μg/dL whereas males only increase by 0.43 μg/dL (i.e. 20-fold higher cortisol response in females). Osmolality change (mmol/kg) was also affected by sex but to a lesser extent than was cortisol. More specifically, the identified QTL explained 14% of the trait variance and sex explained 14.6%. For this QTL, both sexes showed suggestive additive effects with the heterozygote having a value in between the two homozygotes (Table S2). Chloride change (mmol/L) was not sex-dependent, and the identified QTL on BC38 explained 18% of the trait variation. Chloride reduced in the heterozygote individuals by 2.58 mmol/L whereas it stayed approximately the same in the homozygote (0.3 mmol/L; Table 2). Resting blood hematocrit was affected by sex, and two QTL were identified at the chromosome-wide level (23 cM on BC04 and 14 cM on BC25). Together with the sex covariate, these explained 43% of the trait variation, with each QTL explaining approximately 12%. Together, these results indicate the importance of including sex as covariate in these models. These markers will provide targets for selective breeding.

## DISCUSSION

Salmonids are a model system for studying the effects of whole genome duplication (WGD) on genome evolution, sex determination and speciation. Specifically, the evolution of sex determination after WGD and its interaction with heterochiasmy remain active areas of research. In this study we identify the sex-linked chromosome and strong heterochiasmy using the high-density genetic map of Brook Charr. Female recombination rates were 2.7-fold higher than those in the male, and male recombination was highly biased to chromosome ends. When considering only chromosomes expected to be rediploidized this female:male heterochiasmy ratio was 1.85, a lower ratio due to missing regions of the Brook Charr genetic map in residually tetraploid regions. Using recently established correspondence among salmonid chromosomes, we show that the Brook Charr sex chromosome is not the same chromosome arm linked to sex in any other species characterized with high-density genetic maps. However, this chromosome arm (ancestral 15.1) is contained in a fusion within a triple chromosome fused sex chromosome of the congener Arctic Charr. In other salmonid species, some consistencies in sex chromosomes can be viewed, even as distant as Lake Whitefish and members of the genus *Oncorhynchus* (discussed below). We additionally evaluate linkage to 29 reproductive, stress and growth phenotypes (21 not including phenotypes measured at multiple time points) and identify 29 genome-or chromosome-wide QTL on 14 of 42 linkage groups and compare these to known QTL in other salmonids. This work provides markers for selective breeding of Brook Charr as well as insight into the role of heterochiasmy in sex determination and genome evolution in a post-WGD salmonid genome.

### Sex determination post whole genome duplication

In species with genetic sex determination, WGD generates multiple copies of sex chromosomes and this may present challenges to the new lineage (Davidson *et al*. 2009) such as unbalanced gametes and independent segregation of sex chromosomes (Muller 1925) or disruptions in dosage balance in species with heteromorphic sex chromosomes (Orr 1990). However, polyploidization also poses other challenges independent to the duplicated sex chromosome system (Mable 2004; Otto 2007). Nonetheless, ancestral polyploidization resulting in paleopolyploid lineages has occurred throughout plant and animal evolution (Allendorf and Thorgaard 1984; Taylor *et al*. 2003; Session *et al*. 2016), although debate still exists on the effect of polyploidization on diversification (e.g. Clarke *et al*. 2016). Diversification may involve sex determination systems, as paleopolyploids may develop a wide variety of sex determination systems, as observed in the teleosts (Mank and Avise 2009). A wide variety of sex chromosomes are used in different teleost groups including stickleback species (Ross *et al*. 2009; Kirkpatrick 2016) and Medaka (Kondo 2006; Myosho *et al*. 2015).. Ancestral allotetraploids can also develop new sex determination systems, such as the African clawed frog *Xenopus laevis* (ZZ/ZW) having a probable translocated W-specific region (Session *et al*. 2016). However, variable sex determination systems are not unique to paleopolyploid lineages. For example, *X. tropicalis* did not undergo the allotetraploid event but has a unique sex determination system that involves W, Z, and Y chromosomes (Roco *et al*. 2015). The extent of the involvement of polyploidization on the lability of sex determination systems remains to be determined.

Taxa with high rates of turnover in sex chromosomes have indicated that some chromosomes are more likely to become sex chromosomes. This is possibly due to favourable gene content, for example when an autosome contains sexually antagonistic genes it can be repeatedly selected to become a sex chromosome (Marshall Graves and Peichel 2010). A comparative analysis of teleosts indicates repeated independent evolution of the same chromosomes as sex chromosomes throughout evolutionary history (see Table 2 in Marshall Graves and Peichel 2010). Therefore it is not only the master sex determining gene that can be repeatedly utilized by evolution, but also certain chromosomes due to the gene content, which can occur over large evolutionary distance. For example, the teleost tongue sole *Cynoglossus semilaevis* and the chicken *Gallus gallus* (both ZZ/ZW) independently evolved sex chromosomes in homologous chromosomes (Chen *et al*. 2014). Similarly, species from three anuran genera that have diverged for over 210 million years (*Bufo*, *Hyla* and *Rana* spp.) have sex-linked markers that map to the same *X. tropicalis* chromosome with a large region homologous to the avian sex chromosome (Brelsford *et al*. 2013), which the authors suggest is due to independent evolution to the same sex chromosome across the different genera. The platypus *Ornithorhynchus anatinus* has five Y and five X chromosomes, all of which are independent but form a chain at meiosis to co-segregate all together into sperm; this system connects the two sex determination types, with the most degenerate sex chromosome as homologous to the Z chromosome of birds and the least degenerate as that homologous to the X chromosome of mammals (Grützner *et al*. 2004; Charlesworth and Charlesworth 2005). At least three non-homologous sex chromosomes exist within *Xenopus*, and the sex determining region of *X. borealis* shares orthologous genes to mammal sex determination pathways (Furman and Evans 2016). In summary, repeated, independent evolution of the same sex chromosome or use of the same set of specific genes for sex determination therefore has been documented across a variety of animal taxa.

Salmonids have genetically controlled sex determination with XX/XY systems (Thorgaard 1977; Davidson *et al*. 2009). Putative translocation of the sex determining gene to different autosomes has resulted in many different sex chromosomes in different lineages (Woram *et al*. 2003) and even within the same species (Küttner *et al*. 2011; Eisbrenner *et al*. 2013). However, comparing across the phylogeny indicates some noteworthy consistencies. The first consistency is that several species use 3.1 as the sex chromosome, or have this chromosome fused with the sex chromosome, including the fused neo-Y of Sockeye Salmon and the sex chromosome of Coho Salmon (Faber-Hammond *et al*. 2012), as well as the sex-linked LG in Lake Whitefish (Gagnaire *et al*. 2013), identified as 3.1 during map comparisons (Sutherland *et al*. 2016). Relative to the variability seen in sex chromosomes in the salmonids, this is a striking consistency considering that these species have diverged for approximately 50 million years (Crête-Lafrenière *et al*. 2012). This consistency may indicate that either a) 3.1 is an ancestral sex chromosome in the salmonids; or b) the different species converged on this chromosome independently as it contains a gene complement that is highly beneficial to be present as a sex chromosome (Marshall Graves and Peichel 2010; Chen *et al*. 2014; Furman and Evans 2016). The second consistency is that the Brook Charr sex chromosome (15.1) is fused within the sex chromosome of Arctic Charr, but is not the same arm that contains the sex marker in Arctic Charr (Nugent *et al*. 2016), indicating that one of these is a neo-sex chromosome. Furthermore, the middle fused chromosome arm in Arctic Charr is 19.1, which is the newly fused arm in the neo-Y of Sockeye Salmon (Table 1). These observations provide further evidence for the fusion of specific chromosomes together that are beneficial for maintaining within the sex chromosome environment. Finally, intraspecific polymorphism in sex chromosomes occurs in Arctic Charr (Moghadam *et al*. 2007; Küttner *et al*. 2011), and Icelandic Arctic Charr were identified as having a sex chromosome as one of the two homeologs AC01 or AC21 instead of AC04, and state that this is homologous to the sex chromosome of Atlantic Salmon Ssa02 (Küttner *et al*. 2011), which is 9.1 (Sutherland et al 2016), again indicating the potential re-use of the same chromosome as the sex chromosome.

The presence of both the Y chromosome of Brook Charr and the neo-Y of Sockeye Salmon as putative fusions into the neo-Y chromosome of Arctic Charr is worth further investigation because neo-Y chromosomes can influence phenotypic divergence and reproductive isolation, as observed in sympatric Threespine Stickleback *Gasterosteus aculeatus* populations (Kitano *et al*. 2009; Kitano and Peichel 2011). Consistencies across a phylogeny can provide insight into speciation. For example, the Threespine Stickleback and Ninespine Stickleback *Pungitius pungitius* have two different sex chromosomes (LG19 and LG12, respectively), and the Blackspotted Stickleback *G. wheatlandi* has a fused Y-chromosome made up of these two linkage groups (Ross *et al*. 2009), to which the authors suggest multiple independent recruitment of LG12 as the sex or neo-Y chromosome. Furthermore, other sticklebacks have different sex chromosomes (Ross *et al*. 2009). The salmonid diversity and consistencies identified here provide another group for analyzing sex chromosome differences in relation to gene content and speciation, and in salmonids also occurs with the salmonid-specific WGD. As more salmonid genomes are characterized, it will become clearer whether certain sex chromosomes are ancestral or have independently evolved, and whether there is a favourable gene content within often-viewed sex chromosomes.

In the context of sex chromosome fusions and residual tetraploidy, several additional observations on the nature of salmonid sex chromosomes can be made from the present analysis (four genera; eight species; Table 1). The first observation is that sex chromosomes with fusions occur only in species specific fusions rather than conserved fusions in the present data (Sutherland *et al*. 2016). Arctic Charr *S. alpinus* has a sex chromosome that in some individuals involved three fused chromosomes (Nugent *et al*. 2016), and all available evidence suggests this is a species-specific fusion given that these fusions are not present in the more basally diverging Atlantic Salmon nor the congener Brook Charr (Sutherland *et al*. 2016). Y fusions are the most common of sex chromosome fusions (Pennell *et al*. 2015) and can permit other sexually antagonistic genes to be linked to the non-recombining regions (Charlesworth and Charlesworth 1980; Charlesworth *et al*. 2005). Y fusions may also occur due to drift with only slightly deleterious effects (Kirkpatrick 2016), as males have increased fusion prevalence in general and increased repeat content (and thus fusion potential) in degenerating Y (Pennell *et al*. 2015). However, since the same chromosomes that are involved in Y fusions in some species are the sex chromosomes in others (e.g. 15.1 or 19.1, discussed above), it suggests that these fusions could have an adaptive advantage, such as the movement of an autosome with alleles under sexually antagonistic selection into the Y chromosome environment, as discussed by Charlesworth and Charlesworth (1980) and Kirkpatrick (2016). When recombination is low in males (i.e. heterochiasmy), this Y fusion holds an additional chromosome in a constantly lower recombination environment, as it will always be present in males. The use of the same chromosomes as sex chromosomes and as fusion partners within the salmonids merits further study. The second observation is that chromosomes with regions of residual tetraploidy can become sex chromosomes; two of the seven identified sex chromosomes (Chinook Ots17 (23.2) and Atlantic Salmon Ssa02q (9.1)) are chromosomes known to exhibit residual tetraploidy (Brieuc *et al*. 2014; Allendorf *et al*. 2015; Lien *et al*. 2016; Sutherland *et al*. 2016), therefore suggesting that exhibiting residual tetraploidy does not prevent a chromosome from becoming a sex chromosome.

Translocation of a sex determining gene to an autosome and the adoption of the autosome as a new sex chromosome may be possible if the gene moves into linkage with a locus that is under sexually antagonistic selection (van Doorn and Kirkpatrick 2007). The probability of this adoption is increased with the occurrence of an inversion in the region by increased linkage through reduced recombination (van Doorn and Kirkpatrick 2007). The retention still will require that the benefit of the new chromosome is greater than that existing on the original sex chromosome. Interestingly, in the unique sex chromosome of Brook Charr (15.1), there is a large inversion in relation to the other salmonids (Sutherland *et al*. 2016). However, this is an interspecific inversion and has not yet been determined whether it is also heterozygous within the species due to low recombination and resultant challenges of generating male maps. To further characterize this, genome sequence for both the X and Y chromosomes of Brook Charr will be valuable.

The salmonids, being at an early stage of sex chromosome evolution (Phillips and Ihssen 1985) provide a good system to study sex chromosome evolution (van Doorn and Kirkpatrick 2007). As we observed here, reduced recombination occurs consistently in male salmonids, being restricted to the telomeric region opposite the centromere, resulting in a lack of recombination between X and Y in the middle of the chromosome. This may facilitate sex chromosome formation, with tight linkage developing across the entire Y chromosome (Haldane 1922; Nei 1969; Lenormand 2003). Heterochiasmy is not only restricted to the sex chromosome, but rather occurs throughout the genome, as has been viewed in several systems with developing sex chromosomes, such as the European tree frog *Hyla arborea* (Berset-Brandli *et al*. 2008), Medaka (see Kondo *et al*. 2001; Kondo 2006), zebrafish *Danio rerio* (Singer *et al*. 2002), and the salmonids of genera *Oncorhynchus* (Sakamoto *et al*. 2000), *Salmo* (Moen *et al*. 2004) and *Salvelinus* (present study). However, heterochiasmy also occurs in systems with fully developed sex chromosomes, such as humans, where females have ∼1.6-fold higher rates than males, which recombine predominantly at telomeric regions (Broman *et al*. 1998).

Y degeneration can occur from a lack of recombination in sex chromosomes (Charlesworth 1991; Charlesworth *et al*. 2005), and this can also result in degeneration of fused neo-Y, when present. Neo-Y degeneration has occurred rapidly in achiasmate male species such as *Drosophila miranda*, having degenerated after only 1-2 My in the non-recombining state (Steinemann and Steinemann 1998; Charlesworth and Charlesworth 2005). In species with heterochiasmy, even before large degeneration, accumulated substitutions can occur throughout a neo-Y and increased sex-biased gene expression occurs for genes within the neo-Y than the other autosomes, as observed in stickleback (Yoshida *et al*. 2014). These changes are not only degenerative, migration to the Y, and preservation of male-beneficial genes on the Y also occurs, as well as dosage compensation and migration of female-beneficial genes to the X (Bachtrog 2006). Many changes can occur between X and Y when crossovers do not occur throughout the chromosomes.

In the salmonids, sex chromosome turnover by *sdY* translocation may restart the process of Y degeneration (Yano *et al*. 2012b). In species with heterochiasmy rather than achiasmy, occasional crossover between X and Y would also reduce sex chromosome heteromorphism and Y degeneration. This may be the reason for sex chromosomes remaining homomorphic in green toad species (*Bufo viridis*) all which have the same sex chromosomes (Stöck *et al*. 2013), and in several members of the *Hyla* genus of European tree frogs, which also all share the same sex chromosomes (Stöck *et al*. 2011). Regeneration of Y chromosomes by occasional crossover is termed the ‘fountain-of-youth’ hypothesis, and is particularly likely for species with the possibility of sex reversal, as recombination rate is based on phenotypic sex rather than genetic sex (discussed in Perrin 2009). Some salmonid sex chromosomes are heteromorphic (Davidson *et al*. 2009) and accumulate repeats (Devlin *et al*. 1998), this may suggest in some species this regeneration is not occurring. Lack of recombination will be accentuated by inversion accumulation and other differentiation between sex chromosomes reducing meiotic pairing and crossovers. Sex reversal is possible in salmonids (Johnstone *et al*. 1978) and has been observed in the wild, for example in Chinook Salmon (Nagler *et al*. 2001), but the greater extent of this occurring in nature in other salmonids is yet to be determined. Relative effects of sex chromosome turnovers, occasional X/Y crossovers, and large sex chromosomal polymorphisms merits further investigation for which the salmonids are a good model system. The extent of Y or neo-Y degeneration, gene migration, or other aspects of sex chromosome evolution have not yet been explored comparatively in the salmonids. As these aspects may differ among species depending on the length of time the chromosome has been used as the Y chromosome, further investigation into interspecific differences (e.g. 3.1 sex chromosome in both Lake Whitefish and members of *Oncorhynchus*), or intraspecific differences between populations having different sex chromosomes (Eisbrenner *et al*. 2013), will be valuable to determine the history of the sex chromosome evolution in the salmonids.

### QTL mapping, hotspots and consistencies with other species

Knowledge on the genetic architecture of important traits in the salmonids is improving, for example for aquaculture-related traits such as disease resistance (Yáñez *et al*. 2014) and stress tolerance (Rexroad *et al*. 2012), and ecologically-relevant traits such as age-at-sea (Barson *et al*. 2015) and body shape evolution (Laporte *et al*. 2015). In the present study we improve the understanding of genetic architecture of growth, reproductive and stress-response traits by identifying QTL on 14 of the 42 LGs in the Brook Charr linkage map (four fused metacentric and 10 acrocentric chromosomes). This improves the previous analysis of these traits on a low-density map (Sauvage *et al*. 2012a, 2012b) and brings the QTL for these phenotypes into the context of the more characterized high-density Brook Charr map with information on correspondence of arms with other salmonids, probable residual tetraploidy and centromere positions, ancestral chromosomes, and identified sex chromosome. Furthermore, the present analysis was conducted on the female map, which has improved positioning of markers relative to the sex-averaged map or the male map (Sutherland *et al*. 2016), probably as a result of low recombination in the male (as viewed here). Finally, in the present work, phenotypes were investigated for sexual dimorphism and QTL analysis used sex as a covariate when necessary, improving resolution of QTL effects.

Several traits measured in the present study have shown significant heritability in the specific Brook Charr strains used in this study. Stress response as measured by cortisol and glucose responses to transport stress showed mean heritability of 0.60 and 0.61 (± 0.2 SE), respectively (Crespel *et al*. 2011). Often heritability depended on strain, for example heritability of body mass showed mean (± standard error) for domestic, Laval and Rupert strains of 0.61 (± 0.07), 0.37 (± 0.06), and 0.30 (± 0.08), respectively (Crespel *et al*. 2013a). Further, condition factor showed mean heritability (± SE) of 0.09 (± 0.1), 0.032 (± 0.017) and 0.5 (± 0.31) for domestic, Laval and Rupert strains, respectively (Crespel *et al*. 2013b). The different estimates for heritability observed in different strains further indicates the importance of evaluating effects of QTL identified here in multiple strains to identify broader implications of the QTL. Although correlated phenotypes clustered on the map as expected (e.g. length, weight), no clustering was observed for blood and stress-related parameters (i.e. hematocrit, change in cortisol, chloride and osmolality), with each trait having a QTL on a different chromosome. Pleiotropy can occur with both positive and negative genetic correlations between traits with common underlying biology (Mackay *et al*. 2009). This is important to consider in marker-assisted selection, to identify QTL useful for simultaneous selective breeding of multiple traits and to avoid negative correlations between desirable traits (Lv *et al*. 2016). Mapping multiple correlated traits simultaneously can help define regions (Jiang and Zeng 1995). However, it can be difficult to determine whether two traits are truly pleiotropic or whether causal variants for each trait are in in tight linkage, especially when a QTL region is wide (Mackay *et al*. 2009) or when paternal inheritance occurs over long fragments due to low recombination rate (as viewed here).

Consistencies in QTL across multiple species can be useful for identifying regions of the genome with highly conserved roles. Several QTL hotspots have been identified within *Oncorhynchus*, specifically for thermotolerance, length and weight on So6b (Hecht *et al*. 2012), So7a (except weight; also viewed in Rainbow Trout and Chinook Salmon), and So11b (see Larson *et al*. 2015). The corresponding Brook Charr LGs to So6 and So11b (Sutherland *et al*. 2016) did not contain any QTL in the present study, but the corresponding LG to So7a (BC34) contains a length QTL (Table 2). This further implicates this chromosome (ancestral 10.2) as having an evolutionary conserved influence on salmonid growth.

Weight and growth are expected to be highly polygenic traits, therefore requiring many individuals to have sufficient power to identify loci of minor effect (Rockman 2012; Ashton *et al*. 2016). For example, sample sizes of at least 500 individuals may be required to identify QTL accounting for less than 5% of the total phenotypic variance (Mackay 2001). This means that often only large effect QTL are identified, leading to the misconception that these are the norm and to an inflation of the actual percent variance explained by the QTL (Beavis 1997; Xu 2003). A negative relationship between sample size and overestimation of effect sizes occurs in QTL studies of outbred populations (Slate 2013). High powered studies can identify more QTL, such as a recent study in Atlantic Salmon with 1695 offspring and 20 sires, which identified four chromosomes harboring major effect growth QTL (Tsai *et al*. 2015). Similarly, a study in Common Carp *Cyprinus carpio* with 522 offspring and eight families identified 10 genome-wide and 28 chromosome-wide significant QTL for three growth traits, with 30/50 chromosomes containing suggestive QTL (Lv *et al*. 2016). QTL can be detected with fewer individuals, although this may result in overestimation of effect sizes for the QTL (Slate 2013). QTL for polygenic traits growth rate, behavior and morphology were identified in Lake Whitefish with 102 individuals in the mapping family (Gagnaire *et al*. 2013; Laporte *et al*. 2015), and in the present study we identified QTL for many of the traits with 169 or fewer individuals. Since the effect of a QTL can differ in different genetic backgrounds due to epistasis (Mackay 2001), it is therefore important to evaluate the effect of markers in different crosses with different genetic backgrounds to better understand the broader use of the marker (Lv *et al*. 2016), which also gives more confidence on true positives and estimated effect sizes of the QTL (Slate 2013).

The precision of mapping QTL within a family depends on recombination rate (Mackay 2001; Mackay *et al*. 2009). Therefore the low number of crossovers in male salmonids will reduce the overall precision of trait mapping. This effect of heterochiasmy has been used by recent salmonid studies to use a two-stage approach by initially using a sire-based analysis with few markers per chromosome to identify chromosomes of interest followed by a dam-based analysis to more finely resolve the QTL positions (Tsai *et al*. 2015). Heterochiasmy is therefore important to consider when designing QTL experiments for species exhibiting this trait. In the present study, several QTL with very broad regions of elevated LOD were identified (e.g. for length on BC03, BC04, and BC05), which may be due to low recombination and paternally associated haplotypes (see Results). In contrast, many of the other identified QTL in this study have small confidence intervals and high percent variance explained, and therefore will be useful for selective breeding (Table 2; Figure 3).

Although QTL mapping connects nucleotide sequence with trait variation, it generally ignores intermediate phenotypes that can be very useful in determining underlying drivers of traits, and the use of the expression levels of gene transcripts as traits to identify eQTL can provide information on the intermediate steps to generate a phenotype (Mackay *et al*. 2009). Traits queried in eQTL experiments have the additional information on gene location in the genome, providing information on cis or trans-eQTL (Mackay *et al*. 2009). This will be an important next step in determining the underlying causes of the genotype-phenotype interaction in Brook Charr.

## CONCLUSIONS

The relationships between sex chromosomes, heterochiasmy and polyploidization have important influences on genome architecture for key biological traits, but much remains unknown about these interactions. Here we identify the sex-linked chromosome in Brook Charr and compare sex chromosome identities across the salmonids to investigate consistencies. Although many different chromosomes are used as sex chromosomes in salmonids, some consistencies can be identified, even in lineages that have diverged for ∼50 million years, *Coregonus* and *Oncorhynchus*. Sex chromosomes that are contained within fused chromosomes thus far are only observed in species-specific fusions and not in conserved fusions. Heterochiasmy, or differences in recombination rate between sexes, may play an important role in the evolution of sex chromosomes. Heterochiasmy is viewed here in the *Salvelinus* genus, and in other salmonid genera *Oncorhynchus* and *Salmo*, with male recombination less frequent than female, and with male crossovers restricted to telomeric regions. Inversions are also important for sex chromosome evolution, and the Brook Charr sex chromosome from the female map exhibits a large interspecific inversion, although the intraspecific polymorphism of this inversion has not yet been determined. Additional analysis of salmonid genomes is needed to understand the effect of the mobile sex determining gene on phenomena such as Y degeneration. To improve the characterization of important traits and potential for selective breeding, we additionally identify 29 QTL across the genome for growth, reproduction, and stress-response traits, several of which having high PVE and well-refined intervals. Hotspots for multiple traits were not common, but we identify that an earlier identified hotspot in *Oncorhynchus* also contains a length QTL in Brook Charr, further indicating the importance of this chromosome region and the value of identifying orthologous QTL with comparative genomics.

## ACKNOWLEDGEMENTS

This work was funded by a Fonds de Recherche du Québec Nature et Technologies (FRQNT) research grant awarded to Céline Audet, Louis Bernatchez and Nadia Aubin-Horth, a grant from the Société de Recherche et de Développement en Aquaculture Continentale (SORDAC) awarded to Louis Bernatchez and Céline Audet, and a grant from the Spanish Ministry of Education (Grant PR2010-0601) awarded to Ciro Rico. Thanks to G. Côté for laboratory assistance, M. Laporte for discussion on QTL and for comments on the manuscript, M. Lamothe and T. Gosselin for discussion on double recombinants and genotyping errors in RADseq data and to T. Gosselin for exporting the required files from STACKS for QTL analysis. Thanks to C. Nugent for discussion on the sex chromosome of Arctic Charr. Thanks to S. Johnston, an anonymous reviewer, and Editor Ross Houston for providing comments and suggestions that improved the manuscript. During this work, BJGS was supported by an NSERC postdoctoral fellowship, and then an FRQS postdoctoral fellowship.

## SUPPLEMENTAL MATERIAL

**Table S1.** Phenotype average, standard deviation and sample size in males and females. Phenotypes showing any differences between males and females (p ≤ 0.2) included sex as a covariate in the model. Sex-specific phenotypes were only tested within the one sex and therefore had smaller sample sizes.

**Table S2.** Complete QTL table with all identified genome- and chromosome-wide QTLs and associated values, including marker sequence and SNP, and effect size of different genotypes.

**Figure S1.** Heterochiasmy plots for individual chromosomes in the maternal (a) and paternal (b) haplotypes. Grey boxes above the plot indicate probable residually tetraploid chromosome arms, and centromere positions transferred from Chinook Salmon are indicated by stars. When Chinook Salmon chromosomes were not fusions but were metacentrics (n = 2), this is denoted by a ‘?’ to denote the uncertainty as to the centromere position in Brook Charr. However, given the clear pattern of recombination at the opposite end of centromeres in the male, the location for these unknown centromeres is most likely to the opposite side where the recombination events occurred.

**Figure S2.** Correlation plot of phenotypes used in QTL analysis. Phenotype pairs that do not share any individuals for correlation are shown with ‘?’.

**File S1.** Required files for running Rqtl analysis (phenotype (.qua), map (.map) and genotype (.loc)). The map file corresponds to the female map from Sutherland *et al*. 2016. See the Data Availability for code for performing complete analysis with these files.

**File S2.** Fasta file with RAD-seq tags output using the STACKs population module for alleles from all individual offspring and parents.

